# CTCF mediates the cis-regulatory hubs in mouse hearts

**DOI:** 10.1101/2022.11.04.515249

**Authors:** Mick Lee, Loïc Mangnier, Cory C. Padilla, Dominic Paul Lee, Wilson Tan, Wen Hao Zheng, Louis Hanqiang Gan, Ching Kit Chen, Yee Phong Lim, Rina Miao Qin Wang, Peter Yiqing Li, Yonglin Zhu, Steve Bilodeau, Alexandre Bureau, Roger Sik-Yin Foo, Chukwuemeka George Anene-Nzelu

## Abstract

The 3D chromatin architecture establishes a complex network of genes and regulatory elements necessary for transcriptomic regulation in development and disease. This network can be modelled by cis-regulatory hubs (CRH), which underscore the local functional interactions between enhancers and promoter regions and differ from other higher-order chromatin structures such as topologically associated domains (TAD). The Activity-by-contact (ABC) model of enhancer-promoter regulation has been recently used in the identification of these CRHs, but little is known about the role of CTCF on the ABC scores and the consequent impact on CRHs. Here we show that the loss of CTCF leads to a reorganization of the ABC-derived rankings of the putative enhancers in the mouse heart, a global reduction of the total number of CRHs and an increase in the size of the CRHs. Furthermore, CTCF loss leads to a higher percentage of CRHs that cross TAD boundaries. These results provide another layer of evidence to support the importance of CTCF in the formation of regulatory networks necessary for gene regulation.

**Summary:** Deletion of CTCF in mouse cardiomyocytes led to reorganization of the activity-by-contact scores of the heart enhancers and changes in the cis-regulatory hubs

## INTRODUCTION

Unravelling the role of the non-coding regulatory elements in gene regulation is important to gain mechanistic insight into key biological processes such as cell fate determination and disease pathogenesis ^1^ . Accordingly, tremendous effort has been deployed to annotate these regulatory elements in different tissues thanks to innovative sequencing techniques and computational tools ^2-4^. These techniques include Chromatin immunoprecipitation with sequencing (ChIP-seq), Assay for Transposase Accessible Chromatin with sequencing (ATAC-seq), and Chromatin Capture conformation with high-throughput sequencing (Hi-C) ^5,6^. In addition, computational methods such as the Activity by Contact (ABC) algorithm ^3^, and other tools ^7,8^ have been developed to integrate these datasets and identify enhancer-gene (E-G) pairs required for development and disease ^9-11^.

Besides the direct identification of E-G pairs, other studies have demonstrated that enhancers and promoters interact dynamically within a network called cis-regulatory hubs (CRH), which allows for multiple regulatory connections ^12-15^. These CRHs can be defined as 3D regulatory networks that constitute a complex organization of multi-loci connections of enhancers and promoters, connected in 3D space within the nucleus. They highlight direct and indirect contact between genes and distal regulatory elements and can help predict target genes for disease-relevant non-coding SNPs ^14^. They are often bound by a cluster of transcription factors (TF) and have been implicated in gene co-regulation, lineage specification and disease development ^13,14,16^. These hubs differ from other known 3D structures, such as genomic compartments, which constitute euchromatin or heterochromatin regions. They also differ from topologically associated domains (TADs) defined as units within which most enhancer-promoter interactions occur ^17^. Although the identification of these CRHs is often done solely through detailed analysis of 3D contacts from Hi-C data, ^13,18-21^ a recent study proposed building these CRHs using the ABC algorithm ^14^. The ABC algorithm combines ATAC-seq, ChiP-seq and Hi-C to rank the E-G pairs based on their regulatory impact ^3,22^. Thus in addition to identifying and ranking enhancers, the ABC algorithm can be used to generate a more functionally relevant CRH^14^. CRHs built using ABC were strongly enriched for disease-relevant genes and also helped explain disease heritability ^14,23^.

Given that the 3D genome architecture through HI-C is a key component of the ABC algorithm, which is used to construct the CRHs, ^14^, it is of huge importance to understand the role of architectural proteins like CTCF in these CRHs. The CTCF protein often called the “genome weaver” plays a crucial function in the “folding” of the genome and bringing of enhancers into proximity to their distant target genes ^17,24^. We and others have shown the importance of CTCF in regulating cardiac chromatin architecture and heart disease^25,26^, However, CTCF’s role in regulating CRHs has not been explored. In this study, using mouse cardiomyocytes as a model of terminally differentiated non-dividing cells, we employ the ABC model to identify top mCM gene-enhancers and then annotate the CRHs. We analyze the characteristics of the CRHs containing tissue-specific genes, showing that these genes are often found in multi-enhancer CRHs. We also show that in disease, there is a positive regulation of genes and enhancers within a CRH. Finally, we show that loss of CTCF leads to a merging of CRHs and the formation of new CRHs in the *Ctcf*-KO cells with resultant drastic change in the number and distribution of enhancers that cross the ABC threshold.

## RESULTS

### Mouse Cardiomyocyte Enhancer Landscape Identified Through Activity-By-Contact Algorithm

We performed ATAC-seq, H3K27ac ChiP-seq and HiChIP, to generate the ABC scores for the enhancer-gene pairs in the control adult mouse cardiomyocytes. Using an ABC cut-off of 0.01 ^14^, we identified ∼7,000 out of 156,988 putative distal regulatory regions marked by H3K27ac peaks and/or ATAC peaks in the adult mouse CM that crossed the threshold. This represents about 5% of all putative regulatory regions predicted to have strong regulatory effects on their target genes and is similar to what was observed in another study ^3^. With regards to the enhancer-gene (E-G) pairs, globally there were 34,496 E-G interactions with an average of 4.19 connections per gene and 5.29 connections per enhancer in these healthy adult mouse CMs. Figure 1A shows examples of the top enhancers for 2 CM disease-relevant genes *Mybpc3* and *Myh6*. The comprehensive list of ABC-linked enhancers and target genes can be found in supplementary Table 1. A literature search confirmed that at least 3 of the identified enhancers in our study have been validated. These include the *Nppa/Nppb* super-enhancer located upstream of both genes which has been shown to play a critical role in the stress gene response of *Nppa/Nppb* during pressure-overload-induced CM stress ^25,27,28^. A separate enhancer for *Myh7* identified from the ABC scoring has also been validated previously in the mouse heart ^29^. Deletion of this enhancer region led to the downregulation of *Myh7* in mouse hearts^29^. Taken together, these previous studies provide support for the validity of the ABC model.

**Figure 1:**
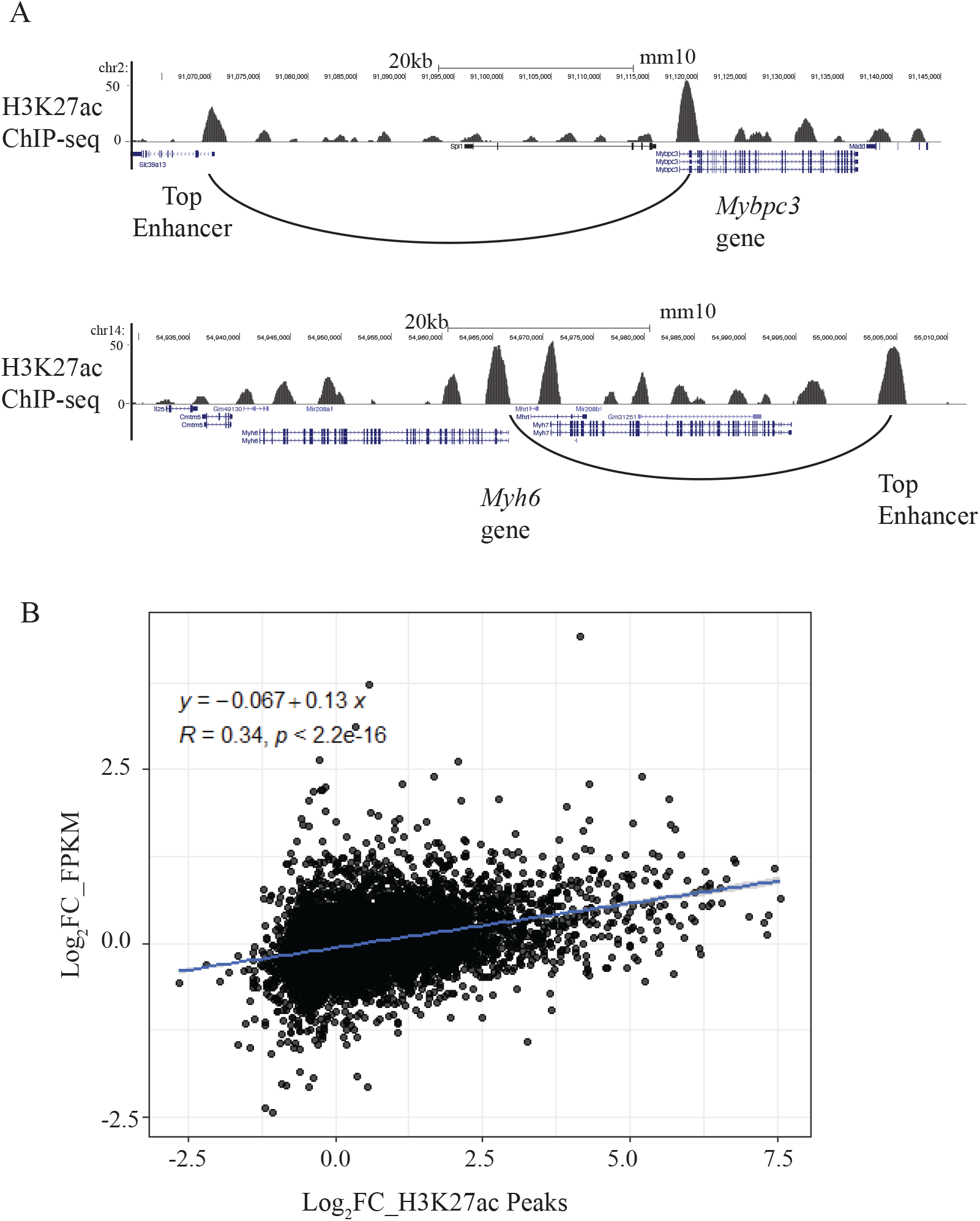
A) USCS screenshot showing 2 examples of CM genes and their top ABC-linked enhancers B) Pearson correlation coefficient showing correlation between differentially expressed genes and differential enhancer peaks for ABC-linked enhancers during cardiac disease

For a better understanding of some of the features of highly ranked enhancers, we focussed our attention on the top 1% and bottom 1% of enhancers based on the ABC score. The enhancers in the top 1% had scores of 0.9 – 1, while those in the bottom 1% had scores of 0.01. First, we analyzed the average distance of these enhancers to the identified target genes. As expected, enhancers with high scores were more likely to be closer to the target genes, with mean and median distances of 83 kb and 54 kb, respectively, as compared to the mean and median distances of the bottom 1% of enhancers which were 950 kb and 699 kb respectively (P value < 0.01). Next, we performed a gene ontology (GO) analysis of the genes linked to these enhancers. The GO biological processes found in the genes linked to top 1% of enhancers include terms such as striated muscle cell differentiation, regulation of blood circulation and actin cytoskeletal organization (Supplementary table 2). These are processes that are important for CM function and include genes such as *Gata4, Mybpc2, Cox7a1* and *Myl3*. On the other hand, the GO for genes in the bottom 1% had generic terms like RNA processing, protein modification and cellular metabolic processes (Supplementary Table 2).

To provide further evidence that these ABC-linked enhancers regulate their putative target genes, we studied the correlation between changes in gene expression versus changes in peak height of the ABC-linked enhancers during pressure overload-induced CM hypertrophy. Using RNA-seq and H3K27ac ChIP-seq datasets from mouse hearts subjected to transverse aortic constriction ^25^, we found that changes in gene expression correlate modestly with changes in H3K27ac peak signals (Pearson correlation= 0.34, *P* < 2.2e-16) (Figure 1B). The correlation was stronger when focusing only on genes and enhancers with greater or less than log_2_ 0.5 fold change after Transverse aortic constriction (Supplementary Figure 1A) (Pearson correlation = 0.51, *P* < 2.2e-16). This result shows that globally, upon external stimuli, gene expression is regulated in the same direction as their ABC-linked enhancers.

### Loss Of CTCF Leads To Changes In The Enhancer Interactome And Changes In The ABC Scores Of Putative Enhancers

Next, to elucidate the role of CTCF in ABC scores and regulation of the CRHs, we also applied the ABC algorithm on the *Ctcf*-KO mouse CMs and observed that 36,226 regulatory elements crossed the 0.01 threshold. This resulted in 71,080 E-G interactions with a similar number of expressed genes (∼8,000) (Supplementary Table 1) in control and *Ctcf*-KO samples. This led to an increase in the average number of connections per gene to 8.14 in *Ctcf*-KO samples. Of the 71,080 enhancer-gene pairs only 20% (14,200 E-G pairs) were shared between control and *Ctcf*-KO. This increase in the number of enhancers with ABC score greater than 0.01 could be due to increased interactions between genes and ectopic enhancers upon CTCF loss. Interestingly we observed that while the highest-ranked ABC enhancer in the control had a score of 0.99, the highest-ranked enhancer in the *Ctcf*-KO sample had a score of 0.78 (scale of 0 – 1). Indeed the average score in the control was 0.054 while the average score in the *Ctcf*-KO was 0.028 (TTest P-value < 2.2e-16). This suggests that the loss of CTCF led to a gain of ectopic enhancers in the *Ctcf*-KO with a redistribution of the ABC scores of the enhancers.

### Loss Of CTCF Alters The Cis-Regulatory Hubs

We integrated the ABC results to generate the CRHs focusing first on the control samples. We identified 1,522 hubs in these control cardiomyocytes with about 70% of the hubs containing fewer than 5 elements (Figure 2A). Figure 2B shows 2 examples of such hubs containing CM genes: *Myom1* a gene involved in the formation of myofibrils ^30^, and found in a simple hub. In contrast, *Tnni3* a CM-specific sarcomeric gene is found in a more complex hub. Consistent with the correlation between gene expression and ABC-linked enhancers in heart disease, we observed that genes and enhancers within the same hubs are positively correlated in the same direction of change during CM stress (Pearson correlation of 0.32 (P <2.2e-16)) (Figure 2C). This suggests a co-regulation of genes and enhancers found within the same hub upon external stimuli. We then ranked CM genes by FPKM and selected the top 50 genes, which were mostly CM-specific genes, to glean biological insights about the characteristics of CRHs that harbour these genes. Our data revealed that highly expressed CM genes were more likely to be in hubs with more regulatory elements than non-highly expressed genes (Figure 2D). Such multi-enhancer hubs have been proposed as a mechanism to buffer against external stresses and genetic perturbations and provide phenotypic robustness to disease-relevant genes ^23,31^.

**Figure 2:**
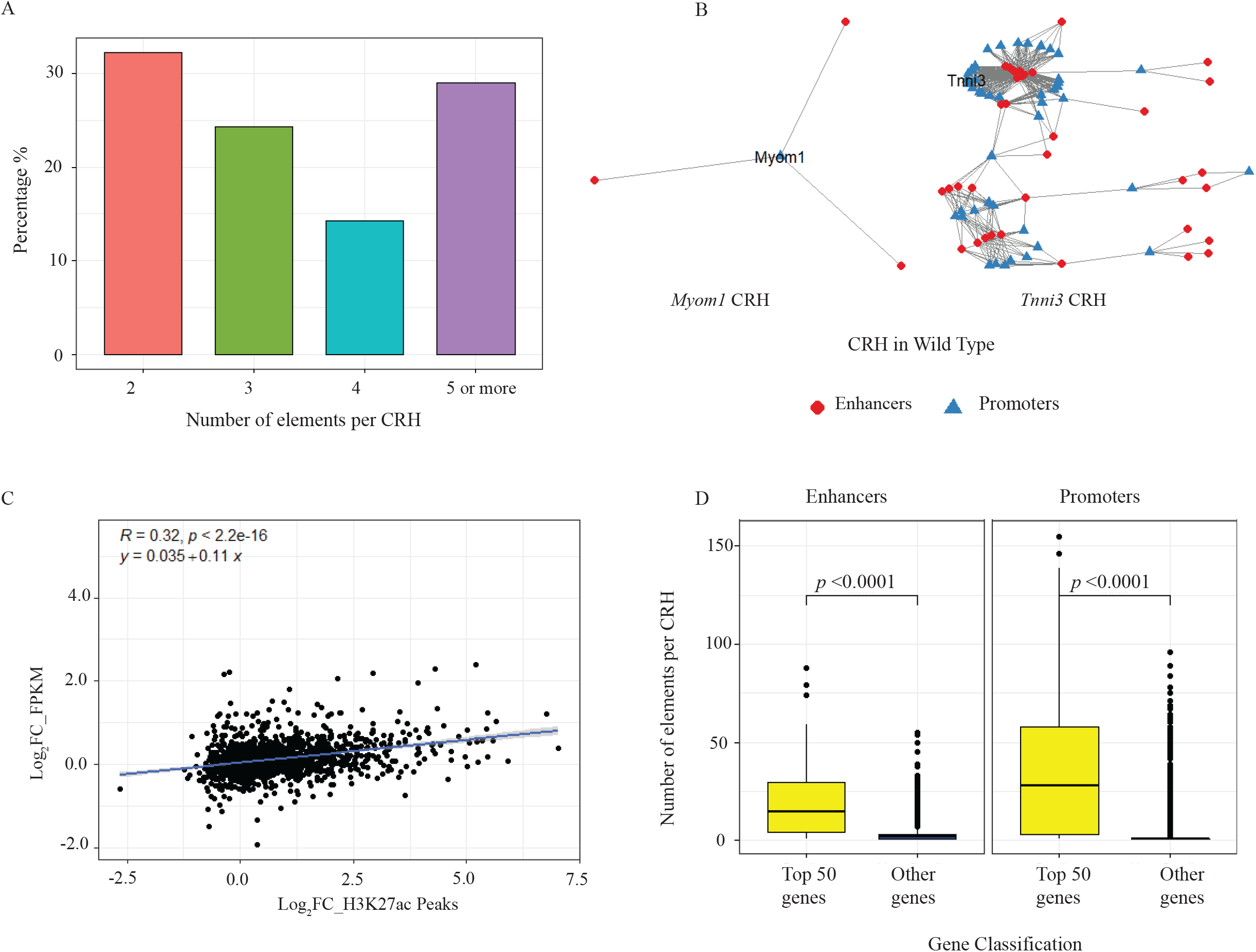
CRH in control Cardiomyocytes A) Bar chart showing distribution of number of elements per CRH B) Examples of 2 CRHs containing CM genes *Myom1* and *Tnni3* in the control cardiomyocytes C) Pearson correlation coefficient between differential expression of genes and enhancer peaks contained within the same CRHs during CM stress D) Box plot showing characteristics of CRHs containing CM-specific genes. These CRHs tend to have more enhancers and promoters

Next, we analyzed the CRHs in the *Ctcf*-KO CMs and found a marked reduction in the total number of CRHs from 1522 in control to 660 CRHs in the KO, accompanied by an increase in the number of elements per hub. Figure 3A shows an example of the same hubs containing 2 CM genes shown in Figure 2A, showing an increase in the number of elements within each hub. Figure 3B shows the distribution of elements per hub, confirming a striking increase in the percentage of hubs with 10 or more elements. Associated with this increase in the percentage of complex hubs, there was an increase in the average number of connections per gene in the *Ctcf*-KO cells, and a decrease in the number of connections per enhancer (Figure 3C, Supplementary Figure 1B). The decrease in the number of connections per enhancer is due to the overall increase in the number of enhancers that crossed the ABC threshold while maintaining the same number of expressed genes. Next, we asked if this reduction in the total number of CRHs in the *Ctcf*-KO was merely due to the merging of CRHs or the formation of new CRHs, and observed both cases. The merging of CRHs was observed primarily in the simple hubs (2-3 elements per hub) located next to each other as 403 (82%) of 492 simple hubs merged with 1 or more other hubs to form larger hubs. For the larger hubs, there was redistribution and formation of new hubs, as genes gained new interactions and lost other interactions. To further confirm the importance of CTCF in the organization of these hubs, we compared CTCF binding sites in the control hubs vs *Ctcf*-KO hubs. Our analysis showed an enrichment of CTCF in the control hubs (fisher exact test odds ratio of 1.69, two-sided P < 2e-16). Next, we analyzed the relationship between CRHs and TADs since TAD boundaries are enriched for CTCF ^25^, and earlier studies have shown that CRHs are generally constrained within TAD boundaries ^14^. Indeed, using the TADs in the control as a reference, our data showed that about 80% of CRHs in the control were contained within the same TAD while only 20% spanned more than 1 TAD. In contrast, about 45% of CRHs in KO were contained within a TAD while 50% of CRHs in the KO spanned more than 1 TAD (Figure 3D) while 5% were not within TAD boundaries, showing that the loss of CTCF leads to a reorganization of the CRHs with the formation of cross-TAD enhancer-promoter interactions.

**Figure 3:**
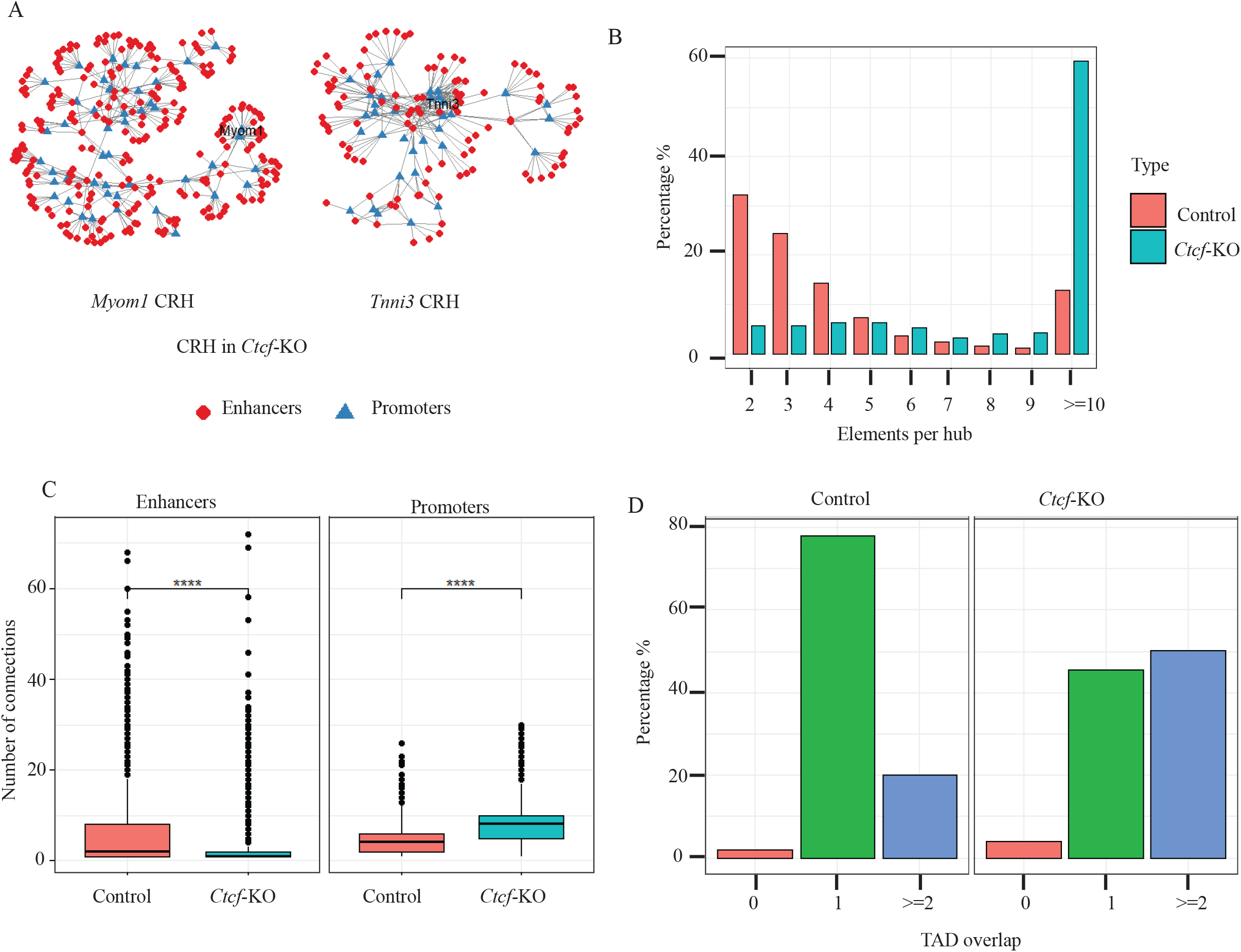
CRHs in the KO cells A) Examples of 2 CRHs containing CM genes *Myom1* and *Tnni3* in the *Ctcf*-KO cardiomyocytes. Compared to the same CRHs in control, there is an increase in the number of elements in each hub B) Bar chart showing distribution of elements per CRH in Control and *Ctcf*-KO. There is an increase in the percentage of CRHs with 10 or more elements from 15% in the Control to 60% in the CTCF KO cardiomyocytes C) Bar chart showing number of connections per gene promoters and per enhancers. There is an increase in the average number of connections per gene in the *Ctcf*-KO CMs. P < 0.01 D) Bar chart showing overlap of CRHs with TADs, 80% of TADs are contained in one TAD in the control while only 40% of CRHs are contained within one TAD in the CTCF KO. In contrast 20% of CRHs span 2 or more TADs in the control, while 50% of CRHs span 2 or more TADs in CTCF KO.

Finally, we sought to identify if CRH changes were directly correlated to gene expression after CTCF loss. We have previously shown that CTCF KO leads to CM dysfunction and global transcriptomic changes after 2 weeks ^25^, here we wanted to gain deeper insight into how that relates to CRH. Our analysis showed that despite the changes in the structure of CRHs, these changes were not directly reflected in gene expression. Indeed there was no correlation between gain or loss of CRH connections and gene expression, suggesting that other secondary effects of the CTCF KO may have come into play to regulate the gene expression profile at 2 weeks when the RNA-seq was performed. This has been observed in other studies that showed that the direct effect of CTCF or Cohesin (another architectural protein) on RNA-seq is observed within a few hours of deletion ^32,33^. As time progresses, other secondary factors come into play and obscure the direct effect of these proteins on gene expression.

## DISCUSSION

Non-coding *cis*-regulatory elements (CRE) play crucial roles in development and disease and hold promise for next-generation therapeutic targets. While different studies by us and others have annotated these CREs and CRHs in different cellular models, the role of CTCF in the ABC-derived CRHs has yet to be studied. First, our study applied the ABC algorithm to the mouse heart, generating to the best of our knowledge, the first such ranked enhancer scores in mice hearts. Our findings confirm the positive correlation between genes and their ABC-linked enhancers in cardiac disease and affirm that genes and enhancers within the same hub are co-regulated during pressure overload-induced CM hypertrophy. Integrating ABC to identify CRHs may thus represent another method to analyze the effect of regulatory SNPs on target genes, as the SNP may affect a target gene when they are in the same hub, even when there is no direct link through pair-wise E-G analysis ^14^. Analysis of the CRHs also suggests that tissue-specific genes are more likely to be contained in hubs with high connectivity and rich in distal elements. This increased number of E-P networks provides redundancy and phenotypic robustness while guaranteeing increased transcriptional activity thanks to the multi-enhancer networks.

Loss of CTCF affects the enhancer interactome, particularly impacting the H3K27ac interactions co-bound by CTCF. This leads to E-Ps that span larger distances and gain of new interactions. The loss of CTCF also leads to a re-organization of ABC-selected enhancers with a direct impact on the CRHs, resulting in fewer hubs but more elements per hub. This implies a crucial role for CTCF in determining which enhancers cross the ABC threshold and the organization of the ABC-derived CRH. While this study focussed on the effect of loss of CTCF on overall CRH structure and its relationship with TAD boundaries, future studies will examine individual CRHs and their relationship to CTCF binding to ascertain how loss of CTCF can lead to local dysregulation of CRHs. Indeed, this finding has implications for mutations that affect local CTCF binding at regions that are not TAD boundaries, as it may lead to the merging of 2 or more CRHs and, thus, the establishment of an indirect connection between genes and enhancers within the newly formed hubs. In addition, given that the binding of CTCF to DNA is methylation-sensitive ^34,35^, this also supports findings that conditions that lead to DNA hypermethylation will reduce local or global CTCF binding ^36^. This loss of CTCF binding will thus lead to the loss of insulation at TAD boundaries with a consequential gain of ectopic enhancers and reorganization of the ABC enhancers ^36^.

## CONCLUSION

Our findings demonstrate that CTCF plays a critical role in determining the regulatory strength of enhancers as well as the membership and organization of the CRHs. Given that the CRHs play a role in disease and cell type-specific gene expression, processes that affect CTCF expression and genomic binding will lead to a disorganization of CRHs and a subsequent effect on transcriptional regulation upon external stimuli. Although we didn’t find a correlation between the reorganization of CRHs and gene expression upon the loss of CTCF, this is mostly due to other secondary effects in the regulation of gene expression. We, however, believe that our findings reveal important principles of the role of CTCF in regulating CRHs and ABC scores. A major limitation of our study is that the resolution of 10kb for our HiChIP is not optimal as each 10kb window may contain various enhancers, and hence, our analysis may miss out on some enhancer-gene interactions. In addition, as the various components of the ABC analysis, namely ATAC, ChIP and HiC, may have technical variations, an independent ABC analysis would be valuable to confirm our findings. We, however, believe that our study forms the foundation for future analysis on the role of CTCF in the ABC enhancer scores and as such the relative contribution of enhancers to each gene. Our future studies will delve deeper into these questions.

## METHODS

### Animal Experiments

Animal experiments were performed under a license approved by the Institutional Animal Care and Use Committee (National University of Singapore). *Ctcf*^flox/flox^ mice harboring LoxP sites flanking exons 3 and 12 of the *Ctcf* gene against a C57/bl6 strain background were used as previously published. AAV9-cTnt-eGFP and AAV9-cTnt-Cre-tdTomato vectors encoding codon improved Cre recombinase or eGFP under the control of cardiac troponin T promoter were used for control and *Ctcf* knock-out respectively ^25^. Experiments were performed on adult 6-to 8-week-old mice. Cardiomyocytes were isolated from mouse hearts after 2 weeks of AAV9 injections for control and *Ctcf*-KO mice. Adeno-associated virus 9 (AAV9) virus for the delivery of Cre recombinase was generated as previously described ^25^.

### H3k27ac HiChip Library Prep And Data Analysis

Mouse adult cardiomyocytes were isolated following previously published protocol ^25^. MNase-based HiChIP assay was performed on 2x10^6^ isolated CMs using the Dovetail Genomics HiChIP MNase kit protocol ^37^. Frozen cells were resuspended in 1× PBS and crosslinked with 3⍰mM DSG and 1% formaldehyde. Washed cells were digested with 0.75⍰µl MNase in 100⍰μl of nuclease digest buffer with MgCl_2_. Cells were lysed with 1× RIPA, and clarified lysate was used for ChIP. H3K27ac antibody (Abcam 04729) was used to perform chromatin immunoprecipitation. The Protein A/G bead pulldown, proximity ligation, and libraries were prepared as described in the Dovetail protocol (Dovetail HiChIP MNase Kit). Libraries were sequenced on an Illumina Novaseq (Supplementary table 3 shows the sequencing QC).

HiChIP paired-end reads were aligned with BWA MEM ^38^ (version 0.7.17r1198-dirty) with the -5SP flag enabled with an index containing only the canonical chromosomes of the mm10 genome (available from the UCSC genome). The resulting alignments were then parsed with pairtools (versions 0.3.0) with the following options --min-mapq 40 –walks-policy 5unque –max-inter-align gap 30 and the thre –chroms-path file corresponding to the size of the chromosomes used for the alignment index. Parse pairs were deduplicated, sorted with pairtools. Valid pairs were identified and through pairtools select (pair type = ‘UU’, ‘UR’, ‘RU’, ‘uu’, ‘uU’ and ‘Uu’) and downsampled to the lowest number of valid pairs in a sample (126 million in LACZ) with pairtools sample. Contact matrices in .hic format were generated with juicetools pre function (version 1.22.01).

FitHiChIP ^39^ (version 9.1) was used to identify significant interactions (“loops”) from the valid subsampled pairs at 10kb resolution with the following settings, loop type = all-to-all, coverage = bias correction, merge redundant loops = Yes, background model = FithHiChIP(L), FDR < 0.1, minimum interaction size =20kb, maximum interaction size = 2 mb. Conditionally unique (meaning only occurring in LACZ or CRE) and shared loops were identified with bedtools pairToPair (version 2.28.0) requiring both loop anchors to overlap the same coordinates be flagged as shared. Loop size was assessed by performing a two-sided Wilcoxson’s rank-sum test on the distribution of loop ranges (distance between the two anchors). Loop anchors were then intersected with the LACZ CTCF binding sites to determine the proportion of loops resulting from CTCF presence. P-values for CTCF mediated loops were obtained through a two-sided Fischer’s Exact test. Aggregate Peak Analysis (APA) ^32^ was computed with juicetools APA function. Loop anchors were used to as the sites to aggregate over at 10kb resolution. APA enrichment scores, loop center from the lower-left (LL) corner is shown as both an APA score (ratio of the mean of center to mean of LL) and a Z-score. APA normed output was used to plot the APA matrices.

H3K27ac HiChIP data were summarized and visualized with R, the package eulurr was used for the shared and unique loop Venn diagrams and ggplot2 for loop size and proportion of loops with CTCF binding site at the anchor. Additionally, ggplot2 was used to plot APA matrices with the geom_raster function, with the color scale set to the same min – max limits across all APA plots. At sites of interest, HiChIP loops files (in longrange format) were used to visualize loops along with ChIP-seq coverage and RNA-seq activity in the WashU Epigenome Browser (https://epigenomegateway.wustl.edu/). Supplementary Table 4 lists all the loops

### RNA-seq

RNA was extracted from 3 biological replicates of control and CTCF KO ventricular CMs (1 million cells each). The paired-end libraries were constructed using Tru-seq kits (Illumina) and resulting libraries were sequenced on the Novaseq platform, generating 2×150-bp paired-end reads. The RNA reads were mapped to the mouse genome with Tophat version 2.0.11 with default parameters, and gene count was computed with htseq-count. Differential gene expression analysis was performed with EdgeR ^40^.

### Chip-Seq Experiment

ChIP experiments on isolated CM (1 million cells each) were performed as described previously ^25^. Briefly, CMs were cross-linked with 1% formaldehyde for 10 minutes at room temperature and quenched with glycine (0.125 mol/L final concentration) ^25^. Cells were then washed in ice-cold PBS and pelleted and lysed in FA lysis buffer (10 mmol/L Tris-HCl, pH 8.0, 0.25% Triton X-100, 10 mmol/L EDTA, 0.1 mol/L NaCl). To facilitate cell lysis, the cell pellet was passed through a 27.5-gauge needle gently 20 times. Nuclei were pelleted by centrifugation resuspended in sonication buffer, and chromatin was fragmented via sonication to an average size of 200 to 300 bp (Bioruptor Plus, Diagenode). Chromatin was immunoprecipitated against H3K27ac (Abcam ab4729) or CTCF (EMD Millipore, catalog No. 07–729) overnight. Antibody-chromatin complexes were pulled down with Protein G Dynabeads (Invitrogen, catalog No. 10003D), washed with 0.1% SDS lysis buffer, and eluted with elution buffer (1% SDS, 10 mmol/L EDTA, 50 mmol/L Tris-HCl, pH 8). After cross-link reversal (4 hours of incubation at 65°C) and proteinase K treatment, immunoprecipitated DNA was extracted. ChIP DNA was quantified by fluorometric quantification (Thermo Fisher Scientific, Qubit dsDNA HS assay kit, catalog No. 32851). Library preparation was performed with the New England Biolabs Ultra II Kit according to the manufacturer’s specifications and sequenced on the Illumina NextSeq High platform.

### ATAC-Seq Experiment And Analysis

The ATAC-seq was performed for both control and *Ctcf* KO CMs according to previously published Omni-ATAC protocol ^41^ Briefly, 50,000 adult cardiomyocytes were pelleted at 500 g for 5 minutes. The cells were resuspended in 100 µl ATAC-resuspension buffer containing 0.5% NP40, 0.5 % Tween-20 and 0.01% Digitonin. The cells were left on ice for 5 minutes after which 1 ml of cold ATAC-RSB containing 0.1% Tween 20 was added to wash out the lysis buffer. The nuclei were pelleted at 500 g for 5 minutes, the supernatant was carefully removed and the nuclei were resuspended in 50 µl transposition mixture containing 25 µl of 2x TD buffer, 16.5 µl of PBS, 5 µl nuclease free water, 0.5 µl of 1% digitonin, 0.5 µl of 10% Tween-20 and 2.5 µl of transposase (Illumina Tagment DNA enzyme 1, Catalogue number 20034198). The reaction was incubated at 37°C for 30 minutes in a thermoshaker with 1000 RPM. After the reaction, DNA was extracted using the NEB Monarch® PCR & DNA cleanup kit (Catalogue number T1030L), PCR was performed using the Illumina/Nextera primers after which Ampure XP beads were used for library cleanup. The resulting library was sequenced on Nextseq.

### ChIP-Seq and ATAC-Seq Data Analysis

For both H3K27ac ChIP-Seq and ATAC-Seq, fastq files were aligned to the mm10 reference genome using the Burrows-Wheeler alignment tool in bwa software ^38^. The corresponding bam files were obtained using samtools ^42^, while peak calling was done with macS2 software ^43^. All peaks with a q-value less or equal to 0.1 were considered as statistically significant and used in the Activity-By-Contact Score ^3^ for CRH construction ^14^. Supplementary table 5 lists H3K27ac peaks in Sham and TAC-treated mouse cardiomyocytes, as well as those in control vs *Ctcf*-KO cardiomyocytes.

### ABC-Score And CRH Analysis

The ABC model defines active enhancers based on a quantitative score of DNAse or ATAC-seq, H3K27ac, and normalized Hi-C contact number^3^. This score is computed relative to a background activity over a 5-Mb window around a candidate element. Then, we set the threshold to 0.01; beyond which a candidate element is considered as a distal element. As an extension of the ABC-Score, CRHs ^14^ were defined as bipartite networks between promoters and distal elements (igraph R package) ^44^ TADs were called using the directionality index as previously described. Pearson correlation analysis was performed using R package

## DATA AVAILABILITY

The raw data supporting the conclusion of this article has been deposited in a public repository.

## ACKNOWLEDGEMENTS

We thank Cantata Bio for providing the HiChIP kits. We thank Dr. Charles Joly-Beauparlant and Dr. Tania Cuppens for helping with data analysis and result interpretation

## AUTHORS’ CONTRIBUTION

M.L, D.P.L, W.Z, L.G, C.C.K, Y.L, R.W, C.G.A.N performed the experiments, M.L, L.M, C.P, W.T, S.B, A.B, C.G.A.N provided data analysis. S.B, A.B, R.F, C.G.A.N Provided supervision for experiments and data analysis

## DECLARATIONS

Animal experiments were performed under a license approved by the Institutional Animal Care and Use Committee (National University of Singapore).

Ethics, Consent to Participate, and Consent to Publish declarations: not applicable.

## FIGURE LEGENDS

Supplementary figure 1

A) Pearson analysis showing the correlation between differentially expressed genes and differential enhancer peaks for ABC-linked enhancers during cardiac disease when taking into account only genes and enhancers with a fold change of log_2_ 0.5.

B) A Box plot showing the number of connections per gene promoters and per enhancers in control and *Ctcf*-KO cells.

